# Co-existence of plasmid-mediated *bla*_NDM-1_ and *bla*_NDM-5_ in *Escherichia coli* sequence type 167 and ST101 and their discrimination through restriction digestion

**DOI:** 10.1101/2024.04.15.589586

**Authors:** Amrita Bhattacharjee, Priyanka Basak, Shravani Mitra, Jagannath Sarkar, Shanta Dutta, Sulagna Basu

## Abstract

The concurrent presence of multiple New Delhi metallo-β-lactamase (*bla*_NDM_) variants within an isolate often goes undetected without the use of next-generation sequencing. This study detects and characterizes dual *bla*_NDM_-variants in *Escherichia coli* through Sanger and whole-genome sequencing. Additionally, a rapid identification method utilizing restriction digestion was designed for detecting variants carrying M154L mutation. Antibiotic susceptibility, minimal inhibitory concentration for meropenem and ertapenem, PCR and Sanger sequencing of *bla*_NDM_ along with genome sequencing using Ilumina and Nanopore technology were conducted. Transmissibility and replicon types of *bla*_NDM_-harbouring plasmids were evaluated. Restriction digestion using restriction enzyme, BtsCI was developed to distinguish between *bla*_NDM-1_ and variants possessing M154L mutations. Two isolates belonging to phylogroups A; ST167 and B1; ST101 and resistant to meropenem and ertapenem (≥16mg/L) were recovered from the blood of a neonate and the rectal swab of a pregnant woman respectively. *bla*_NDM_ was detected by PCR, and Sanger sequences of *bla*_NDM_ showed two peaks at 262 (G & T) and 460 (A & C) nucleotide positions indicative of more than one *bla*_NDM_ variant. Hybrid assembly confirmed co-existence of *bla*_NDM-1_ and *bla*_NDM-5_ in each isolate. *bla*_NDM-1_ was located on IncY (ST167) and IncHI1A/HI1B (ST101), while *bla*_NDM-5_ was on a IncFIA/FII (ST167) and IncC (ST101) plasmids. Digestion with BtsC1 could discriminate *bla*_NDM-1_ and *bla*_NDM-5_. Co-existence of multiple *bla*_NDMS_, *bla*_NDM-1_ and *bla*_NDM-5_ in epidemic clones of *E. coli* is concerning. Restriction digestion method and Sanger sequencing can facilitate quick identification of dual *bla*_NDM_ variants in single isolate.

**Importance:** The global dissemination of antimicrobial resistance genes is a serious concern. One such gene, *bla*_NDM_, has spread all across the globe via plasmids. *bla*_NDM_ confers resistance against all β-lactam antibiotics except monobactams. Most of the earlier literature reported presence of single *bla*_NDM_ variants. However, this study reports the prevalence of dual *bla*_NDM_ variants (*bla*_NDM-1_ and *bla*_NDM-5_) located on two separate plasmids identified in two distinct *E. coli* epidemic clones ST167 and ST101; isolated from a septicaemic neonate and a pregnant mother respectively. *bla*_NDM-5_ differs from *bla*_NDM-1_ due to presence of two point mutations i.e., V88L and M154L. This study detected dual *bla*_NDM_-variants through Sanger sequences and further validated through hybrid-genome assembly. Detection of multiple *bla*_NDM_-variants in a single isolate remains difficult until genome sequencing or southern blotting are carried out. Hence, a simple restriction digestion method was devised for rapid screening of dual *bla*_NDM-_variants containing M154L mutation.

## Introduction

New Delhi metallo-β-lactamase (*bla*_NDM_) is the fastest and widest spread carbapenemase that has triggered an alarming threat worldwide since its identification in 2009 (1, 2). Around 60 variants of *bla*_NDM_ have been reported worldwide of which *bla*_NDM-1_, *bla*_NDM-5_ and *bla*_NDM-7_ are prevalent (3). NDM confers resistance against all β-lactam antibiotics including carbapenems, the drug of last resort (4).

Most studies report presence of single *bla*_NDM_ variant in a bacterial genome with few exceptions where two copies of *bla*_NDM_ variants (*bla*_NDM-1/NDM-5_) were present either in chromosome or plasmids in a single bacteria such as *Pseudomonas aeruginosa, Escherichia coli, Klebsiella michiganensis* and *Acinetobacter johnsonii* (Table S1). *E. coli*, being a commensal and an opportunistic pathogen acts as a reservoir of acquired antimicrobial resistance determinants eventually transferring it to other species (5).

In this study, we identified two *E. coli* isolates from blood (neonate) and rectum (adult), possessing two different variants of *bla*_NDM_. To the best of our knowledge, this is the first report where carriage of bi-variant *bla*_NDM_ in *E. coli* are reported along with their precise characterization and genome analysis.

## Results and discussions

EN5349 was isolated from blood of a septicaemic neonate whereas IN-MR210EC from rectal swab of a pregnant mother hospitalized for delivery [as part of collaborative study named, Burden of antibiotic resistance in neonates from developing societies (BARNARDS)]. The mother and the baby were not paired. Both isolates were belonged to phylogroups A; ST167 (EN5349) and B1; ST101 (IN-MR210EC) which are epidemic clones; conferring resistance to carbapenems and other antibiotics, susceptible to tigecycline and colistin. Serotypes, plasmid types, pMLST, *gyrA* and *parC* mutations, prevalence of resistance and virulence determinants are presented in Table 1.

**Table 1.**
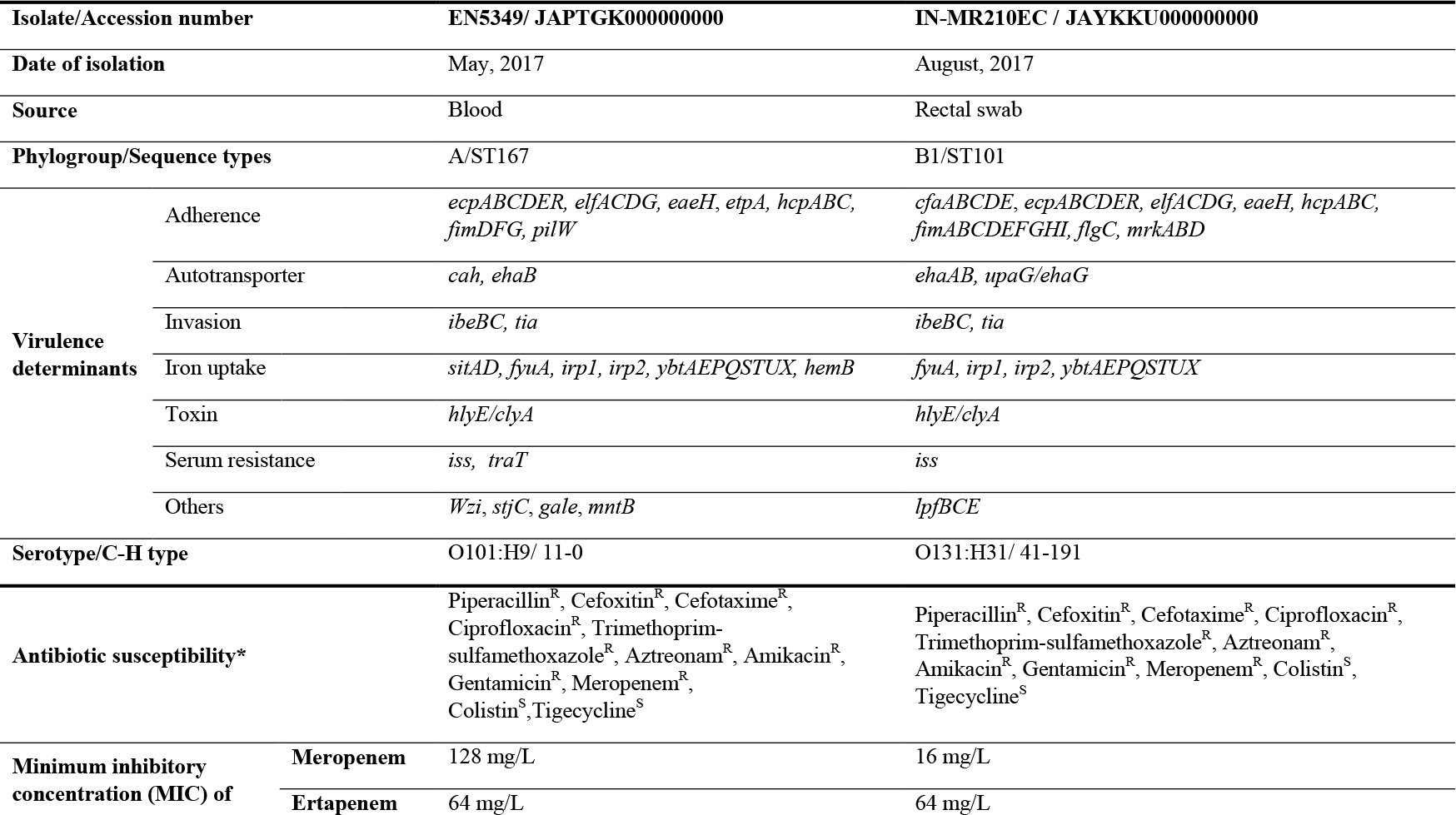
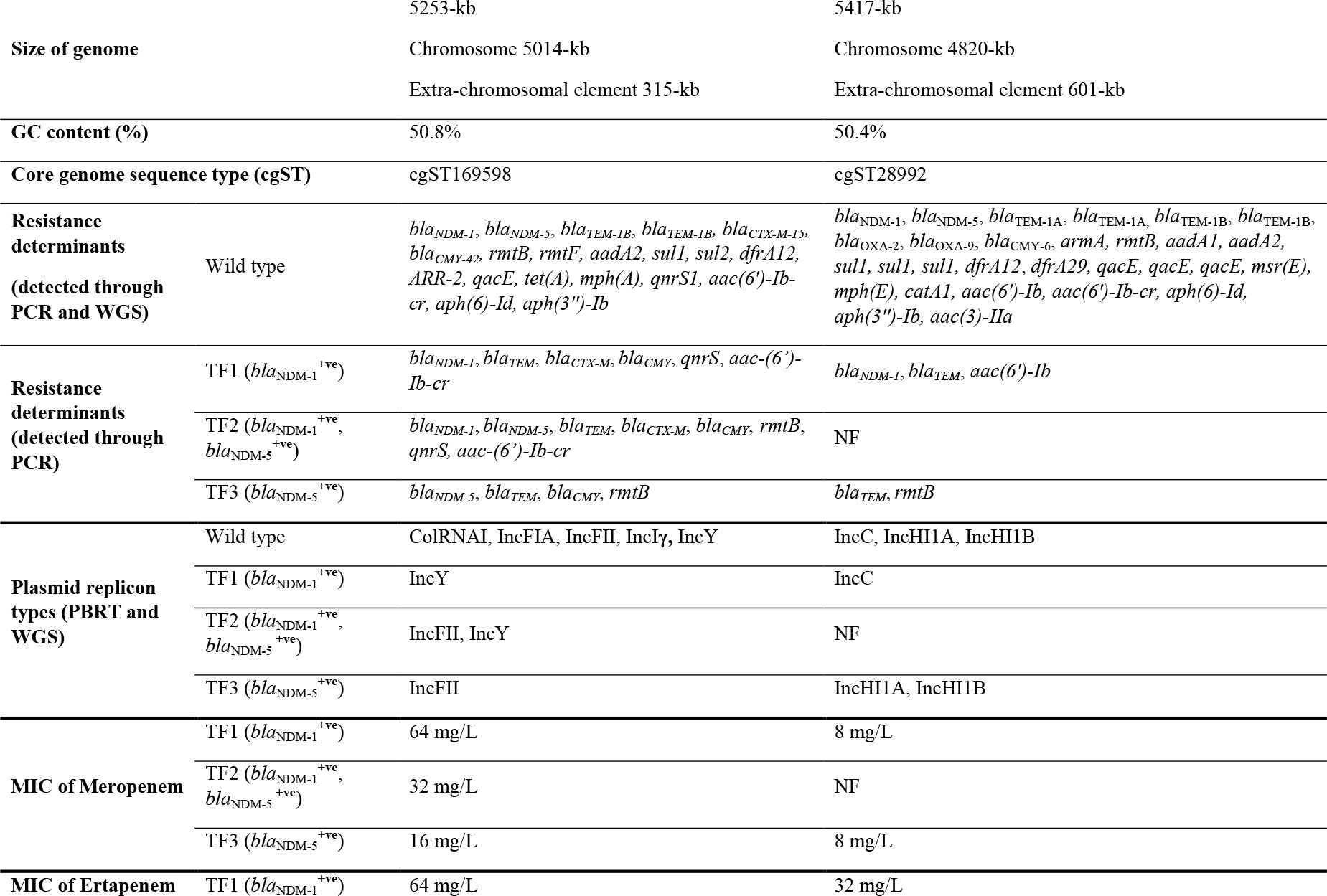
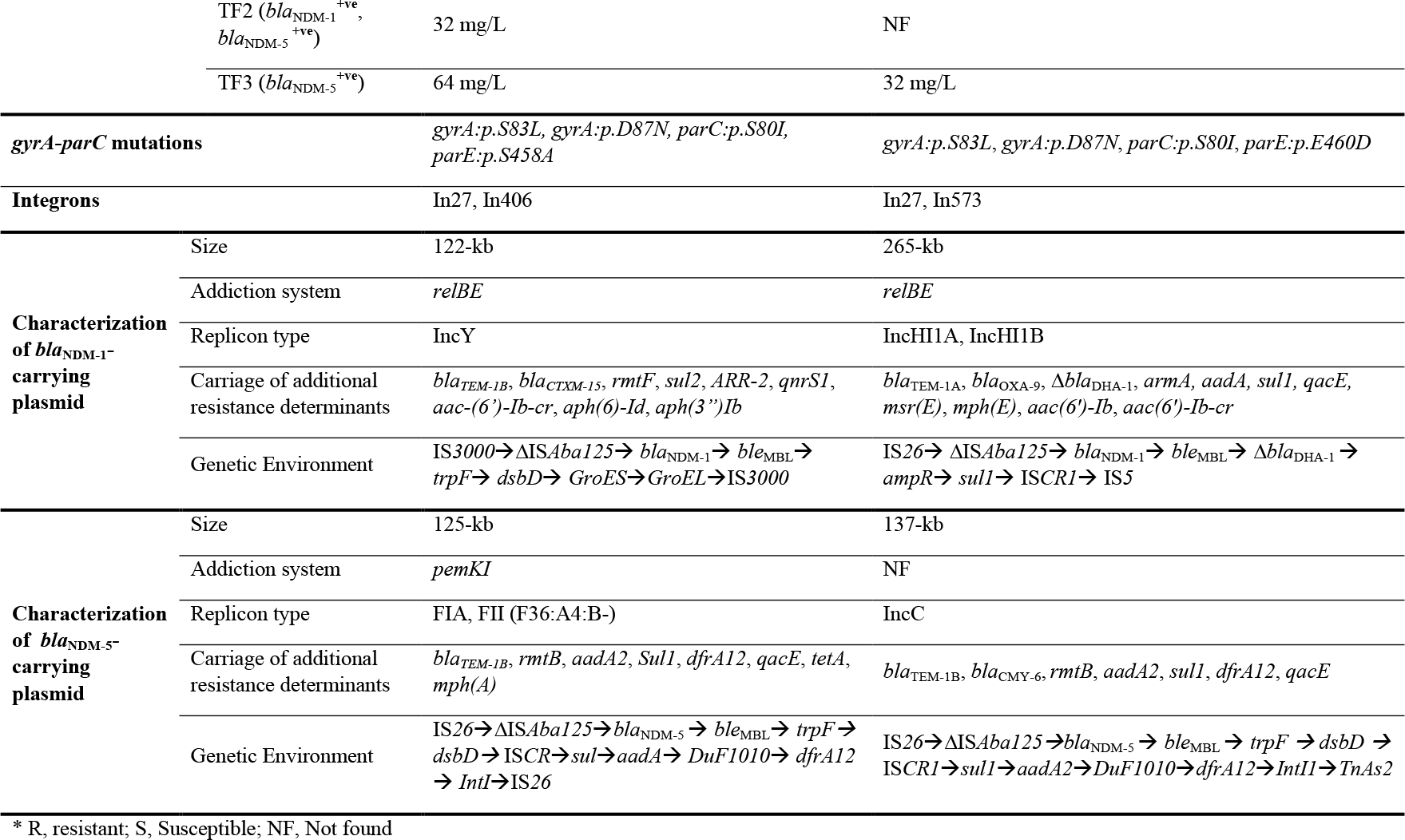
Genome-based characterization of EN5349 and IN-MR210EC along with characterization of transformants (TFs) in terms of resistance determinants and plasmids.

Presence of *bla*_NDM_ in isolates was confirmed by Sanger sequencing. Chromatogram of sequences (forward and reverse) for both isolates depicted presence of two sharp peaks at 262 (G & T) and 460 (A & C) positions. Two different base calls at a single position suggested amplification of more than one *bla*_NDM_ in a single isolate (Fig. S1). G at position 262 and A at 460 correspond to sequence of *bla*_NDM-1,_ whereas G262⟶T (V88L) and A460⟶C (M154L) matched with *bla*_NDM-5_ (6). The meticulous checking of chromatogram helped the detection of dual *bla*_NDM_ copies.

Hybrid assembly revealed that both isolates were carrying two copies of *bla*_NDM_ (*bla*_NDM-1_ and *bla*_NDM-5_) along with two copies of *bla*_TEM-1B_ which may increase enzyme production, and also compensate for the loss of any one plasmid. Additionally, IN-MR210EC carried multiple copies of other resistance determinants (Table 1). *bla*_NDM-1_ was harboured in IncY [P1-EN5349; 122-kb], and IncHI1A/HI1B [P1-IN-MR210EC; 265-kb]; whereas *bla*_NDM-5_ was present in IncFIA/FII [P2-EN5349; 125-kb] and IncC [P2-IN-MR210EC; 137-kb], respectively. The genetic environment of *bla*_NDM-1_ comprised of Tn3-like IS*3000* and IS*26*-like family transposase upstream of EN5349 and IN-MR210EC respectively along with truncated IS*Aba125* in both cases but the downstream regions varied (Table 1). However, in case of *bla*_NDM-5_, IS*26*-like family transposases and a truncated IS*Aba125* were found upstream and *ble*_MBL_ downstream followed by *trpF*, *DsbD*, IS91-like IS*CR1* family transposase for both isolates (Table 1 and Fig. S2). Although the plasmid replicon types and plasmid backbones were different, similar genetic context may suggest the transposition of *bla*_NDM-5_. Different determinants such as, plasmid replication initiation factors (*repA, repB*) (7); conjugation/type IV secretion system (T4SS) (*virB, virB9, virB11, virB6, virB4*, Rhs) (8); partitioning (*parA, parB, parM*) (9), stable inheritance (*pemK, pemI*) (10); site specific recombinase (*xerC*) (11) and transfer/mobilization (*tra*-operon system, *mob*I) (Fig. 1 and Fig. S2) were present.

**Fig 1.**
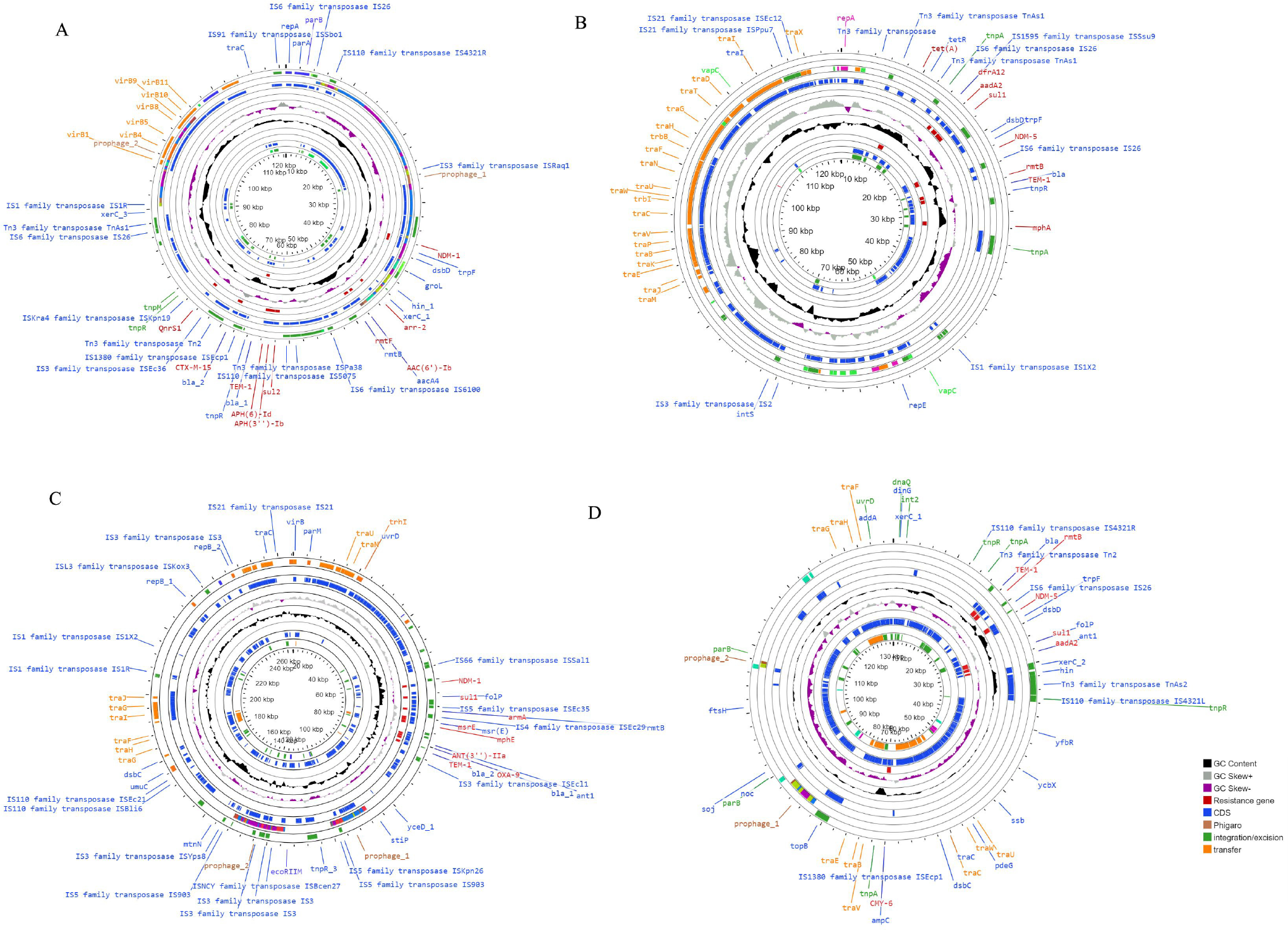
Circular map of the *bla*_NDM-1_ and *bla*_NDM-5_-harbouring plasmids in different *Escherichia coli*, ST167 and ST101. Circular graphs represent the genetic context of *bla*_NDM-1_ in IncY (A) & *bla*_NDM-5_ in IncFIA, IncFII (B) in EN5349 (ST167) isolated from blood and *bla*_NDM-1_ in IncHI1A, IncHI1B (C) & *bla*_NDM-5_ in IncC (D) in IN-MR210EC isolated from gut. The inner circle is depicted as GC content in black and the outer circle represents GC skew in grey and purple. Resistance gene is denoted in red, coding sequences (CDS) in blue, prophage regions (phigaro) in brown, genes involved in transfer is in orange and integration/excision in green.

Randomly selected TFs possessed either *bla*_NDM-1_ or *bla*_NDM-5_. Only few TFs of EN5349 (EN5349.TF3) co-harboured *bla*_NDM-1_ and *bla*_NDM-5_ as confirmed by Sanger Sequencing. WT and TFs exhibited high MIC values for meropenem and ertapenem (Table 1) that may lead to treatment failures particularly when other resistance determinants are present along with *bla*_NDM_.

Study plasmids showed close resemblance with some globally-reported plasmids, in different species of Enterobacterales. P2-EN5349 [pMLST F36:A4:B-] showed similarity with *bla*_NDM-5-_possesing plasmids isolated mostly from *E. coli* reported from Canada, Czech Republic, Switzerland, Rome, Myanmar, Bangladesh, China, Thailand and India. P2-IN-MR210EC (*bla*_NDM-5_-possessing) showed similarity with an environmental sample of *E. coli*, reported from Switzerland. *bla*_NDM-1_ harbouring study plasmids exhibited similarities with plasmids reported from India and Thailand (Fig. 2 and Table S2).

**Fig 2.**
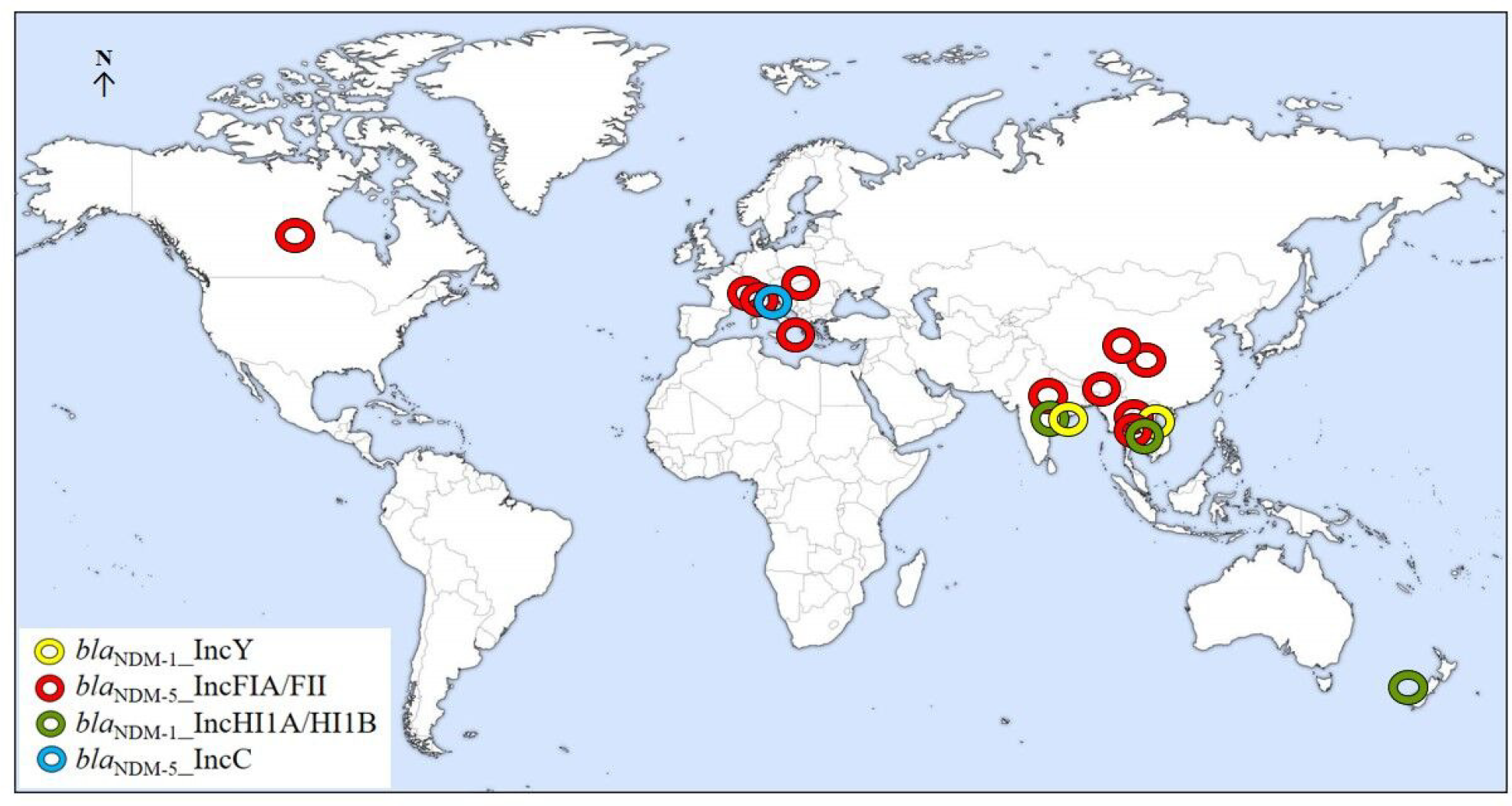
Worldwide prevalence of plasmids similar to study plasmids. Globally reported plasmids similar (≥99% nucleotide identity) to study plasmids are denoted in four different colours such as yellow (*bla*_NDM-1_ in IncY), red (*bla*_NDM-5_ in IncFIA/FII), green (*bla*_NDM-1_ in IncHI1A/HI1B) and sky blue (*bla*_NDM-5_ in IncC). P2-EN5349 [pMLST F36:A4:B-] showed similarity with *bla*_NDM-5_-possesing plasmids reported mostly in *E. coli*; from several countries such as Canada, Czech Republic, Switzerland, Rome, Myanmar, Bangladesh, China, Thailand and also from India (Accession NZ_CP033159.1). In contrast, P2-IN-MR210EC (*bla*_NDM-5_-possesing) was found similar to only one plasmid harbored in *E. coli* isolated from environmental sample reported from Switzerland (Accession NZ_CP048374.1). *bla*_NDM-1_ harboring study plasmids showed identity with plasmids reported from India and neighboring country Thailand.

Dual *bla*_NDM_ variants in a single isolate may be confirmed by southern blotting and hybrid assembly which are expensive, time consuming and difficult to execute for a diagnostic laboratory. Hence, a novel restriction digestion based method was introduced to distinguish between *bla*_NDM_ variants (NDM-4,5,7,8,12,13,15,16b,17,19,20,21,27,35,36,37) with or without M154L mutation. *bla*_NDM-1_ has two recognition sites for BtsCI (Fig. S3) which generates three DNA fragments (255, 215 and 355 bp). Presence of a mutation (A460⟶C) corresponding to M154L alters the second recognition site (GG<u>A</u>TG⟶GG<u>C</u>TG) which BtsCI is unable to cleave, resulting in two fragments of 255 bp and 570 bp (Fig. S3). Hence, most M154L possessing variants (including *bla*_NDM-5_) will produce fragments different from *bla*_NDM-1_ variant. Since study isolate possessed both *bla*_NDM-1_ and *bla*_NDM-5_, 4 DNA fragments such as 215 bp, 255 bp, 355 bp and 570 bp (Figure S4) were generated which confirmed the presence of *bla*_NDM-1_ along with a variant possessing M154L mutation (4).

## Conclusion

Our study adds to the very few studies that reported more than one variant of *bla*_NDM_. The presence of multiple *bla*_NDM_ variants in *E. coli* collected from a septicaemic neonate and a pregnant mother (not paired) is worrisome. Emergence and spread of such organisms in India is of immense public health consequence. Two different variants of *bla*_NDM_ in different plasmids and their stability in antibiotic-loaded environment calls for timely identification.

## Materials and methods

Two clinical *E. coli* isolates, EN5349 and IN-MR210EC were assessed for antibiotic susceptibility by disk diffusion assay and minimal inhibitory concentration (MIC) for meropenem, ertapenem and colistin (Sigma-Aldrich, Steinheim, Germany) were determined by broth-micro dilution method (12). PCR amplicons of *bla*_NDM_ were sequenced using primer pairs (Table S3) in Applied Biosystems, DNA Analyzer (Perkin Elmer, USA). Both short-read (Illumina NextSeq 500 platform, San Diego, CA) and long-read based (Oxford Nanopore, UK) sequencing technologies were used for genome sequencing (13). Unicycler was used to generate hybrid assemblies which were further used for downstream analysis (supplementary methods)(14). From hybrid assemblies, *bla*_NDM_ harbouring plasmids were constructed and were searched in the bacterial plasmid database (PLSDB) (15) for similar complete plasmid sequences. Plasmids showing nucleotide identity (≥99%), same replicon types with similar *bla*_NDM-_variants with reference to study plasmids were compared.

Transfer of *bla*_NDM_ into *E. coli* J53 Az^r^ strains (100mg/L) was attempted by solid-mating conjugation assay. Electro-transformation was achieved using plasmids (PureYield™ Plasmid Midiprep System, Promega, United States) and transformed to recipient *E. coli* DH10B cells (Invitrogen, CA, USA)(12, 13). Transformants (TFs) were selected on LB agar (Difco™, Lennox) supplemented with 2mg/L meropenem (Sigma-Aldrich, USA) and presence of *bla*_NDM_ was confirmed. MIC values were checked for meropenem and ertapenem for TFs. Plasmid replicon types were determined by PCR-based replicon typing (PBRT, Diatheva, Italy) for wild-type isolates (WTs) and TFs (12). PCR amplicons of *bla*_NDM_ gene were digested for 4 hours using the BtsCI enzyme at 50ºC.

## Data availability

Genome data was submitted to the NCBI database with accession numbers **JAPTGK000000000** and **JAYKKU000000000** (Table S1).

## Ethical approval

This study was conducted on archived strains, hence ethical approval was not required.

## Acknowledgement

We would like to express our gratitude to Thomas Jove (INTEGRALL) for helping in curation of integron sequences. We thank Dr. Wriddhiman Ghosh for his valuable inputs to execute the hybrid assemblies.

We sincerely thank all the staff and clinicians of Department of Neonatology, SSKM and IPGME&R hospital for their co-operation in handling and sending the samples.

## Transparency declarations

None to declare.

## Author Contributions

SB contributed to the conception of the work, analysis, interpretation of data, and drafting of the manuscript. AB performed the all experimental work and contributed to the acquisition of laboratory and WGS data with manuscript drafting. PB and JS performed the WGS data analysis with manuscript editing. SM contributed to the analysis of data. SD contributed to review of data. The final manuscript was read and approved by all authors.

